# Innovative, rapid, high-throughput method for drug repurposing in a pandemic – *a case study of SARS-CoV-2 and COVID-19*

**DOI:** 10.1101/2022.12.25.521651

**Authors:** Shaibu Oricha Bello, Abdulmajeed Yunusa, Adamu Ahmed Adamu, Mustapha Umar Imam, Muhammad Bashir Bello, Abdulmalik Shuaibu, Ehimario Uche Igumbor, Zaiyad Garba Habib, Mustapha Ayodele Popoola, Chinwe Lucia Ochu, Aishatu Yahaya Bello, Yusuf Yahaya Deeni, Ifeoma Okoye

## Abstract

Several efforts to repurpose drugs for COVID-19 treatment have largely either failed to identify a suitable agent or agents identified did not translate to clinical use; either because of demonstrated lack of clinical efficacy in trials, inappropriate dose requirements and probably use of inappropriate pre-clinical laboratory surrogates of effectiveness. In this study, we used an innovative algorithm, that incorporates dissemination and implementation considerations, to identify potential drugs for COVID-19 using iterative computational and wet laboratory methods that highlight inhibition of viral induced cytopathic effect (CPE) as a laboratory surrogate of effectiveness. Erythromycin, pyridoxine, folic acid and retapamulin were found to inhibit SARS-CoV-2 induced CPE in Vero cells at concentrations that are clinically achievable. Additional studies may be required to further characterize the inhibitions of CPE and the possible mechanisms.

**Funding:** TETFund Covid-19 Special Intervention Research grant(grant number TETFund/DR&D/CE/ SI/COVID-19/UDUS/VOL 1)

## 1. Introduction

Coronavirus disease 2019 (COVID-19) pandemic was not a rude surprise because impending pandemics had long been predicted based on scientific data, and preparations existed for early detection and response^1^. When SARS-CoV-2 finally crossed the species barrier and initiated a rapidly evolving worldwide morbidity and mortality events, even the most advanced health systems in terms of resources and scalability were easily overwhelmed^2^. Nonetheless, the rapid discovery, dissemination and implementation of effective vaccine is estimated to have saved millions of lives^3^ and can largely be traced to the availability of repurposed mRNA technology^4^ initially developed with therapeutics in view, though other vaccine platforms were also successful^5^. A major drawback was the lack of equity in vaccine distribution with developed countries acquiring vaccine doses in multiples of their population needs while less developed economies that had neither the resources for competitive purchases nor capacity for unassisted vaccination campaigns were late in vaccine deployment^6,7^. Efforts at identifying therapeutics for people who do develop COVID-19 have had only limited success with most early enthusiasm at discovery of effective repurposed drugs rapidly and consistently failing randomized clinical trials^8^. With the exception of Dexamethasone and related drugs, the few effective COVID-19 therapeutics like Nirmatrelvir/ritonavir (Paxlovid) and even less effective Remdesivir (Veklury) were priced out of the reach of most countries, especially developing economies^9^. It is informative that both Nirmatrelvir and Remdesivir are essentially repurposed^10^.

Repurposing drugs appears to be an effective approach to rapidly discover therapeutics for emerging infectious diseases. What seems to be faulty is the lack of factoring dissemination and implementation into the initial repurposing inquiry. Such inclusion would involve reflection on potential hindrances to rapid worldwide use of the drug such as cost, inherent toxicity^11^, worldwide accessibility, ease of scalability and other factors that may together be described as *off-label use likelihood* (e.g., familiarity of clinicians with the drugs, background acceptability by the population, among others). Also, most of the effort at repurposing drugs have been conservative on molecular targets, by frequently restricting screening to inhibition of one target in the virus and /or disease evolution pathway^12,13^. However, lessons from HIV suggests that therapeutics with multiple targets may have higher effectiveness and tremendously lower the probability of resistance by the virus^14^. It has been suggested that a new approach to drug repurposing is needed^10^ and that for emerging viral diseases, combination therapy or targets should be prioritized^15^. This so called one-drug-multiple-target approach is attractive and has been used successfully for non-viral diseases^16^. Fortunately, current in-silico technology and supporting ecosystem (databases) enable highly cost-effective screening of multiple small molecules for potential activity at multiple targets and greatly reduces wet laboratory transition time and cost^17^.

Methods of wet laboratory validation frequently used is inhibition of viral multiplication as measured by reduced viral number on quantitative PCR (qPCR)^18^. However, qPCR is not optimized for rapid high throughput screening of large number of compounds. Furthermore, reduction in viral load may not be predictive of clinical efficacy, especially where the mechanism of viral disease involves cellular cytopathy and a low viral threshold for cytopathy exists^19^. SARS-CoV-2 induces cellular cytopathy in its target cells and inhibition of cytopathic effect has been employed as a high throughput screening method for drug repurposing against COVID-19^20^.

This study was therefore designed as a cost-effective, easy to conduct, high throughput procedure to repurpose drugs using in silico screening against multiple SARS-COV-2 and COVID-19 targets, then select from the hits using criteria that will enhance dissemination and implantation, then conduct wet laboratory validation of the selected hits using inhibition of cytopathic effect (CPE) as a measure of efficacy.

## 2. Materials and Methods

### 2.1 Theoretical Framework

Although orthodox medicines are developed for narrow indications, by their nature, small molecules (drugs) have the capacity for activity at various sites, some of which may have clinical benefits^21–23^. Drugs with multiple sites of activity are essentially ‘combination therapies’ and therefore may be expected to show a high threshold for viruses to develop resistance. For example, although the rate of resistant mutation is unknown with SARS-CoV-2, it is estimated as 1 in 10^6^ for HIV-1 virus^24–26^. This suggests that a three-drug combination with different sites of action will acquire 1 in 10^18^ resistant viruses: a number much higher than the possible viral loads in the body. When repurposing drugs for viral infections, it is therefore prudent to screen drugs for potential activity at all known targets (confirmed and putative) in the viral cycle and select those drugs with the highest overall score (negative binding score) across the targets. It is also prudent that the minimum potential sites of activity of a candidate drug for selection will thus be three sites.

Although different virus families interact with their host cells in very different ways, they have historically been categorized as either generating abortive, persistent or cytolytic infections^27^. SARS-CoV-2 is known to cause CPE in infected cells and SARS-CoV-2 Spike variant has been shown to have enhanced cytopathic and fusogenic effects in infected cells^28^. We, therefore, propose that inhibition of virus induced CPE by drugs would be more rapid and predictive of clinical efficacy than inhibition of viral replication, especially where virus induced cellular death is a major pathway of disease, as with COVID-19. This is because viral load suppression alone may not go below the viral threshold for CPE though the suppression may be of multiple folds compared to control. Lack of inhibition of CPE may be one of the reasons many repurposed drugs fail. In addition, inhibition of CPE could be more cost-effective to rapidly screen multiple batches of candidate drugs than qPCR.

Also, it has been shown that only about 12.5% of positive research findings are ever disseminated and implemented (DI) and that even these (DI) take over 10 years^29^. To improve this status, a theory-informed approach built into research design has been suggested^29–32^. We propose that DI of study results are more successful if designed into the research a priori by involving stakeholders like infectious disease clinicians, policy makers and clinical protocol developers in a needs assessment and other phases of the study: thus, making them, by their participation, aware of the rationale and thoroughness of the primary data generation and its analysis.

### 2.2 Repurposing Algorithm

The repurposing algorithm used in this study includes (i) Needs assessment, NA (ii) Evaluation and Efficacy Studies, EES and (iii) Dissemination and Implementation Steps (DIS), which were iterative with continuous review and enrichment as knowledge about SARS-CoV-2 and COVID-19 accumulated (Figure 1). Where required, consensus was reached by modified Delphi protocol^33^ involving subject specialists and measurements were made using 4-points Likert’s scale to evoke forced choices and avoid neutrality.

**Figure 1:**
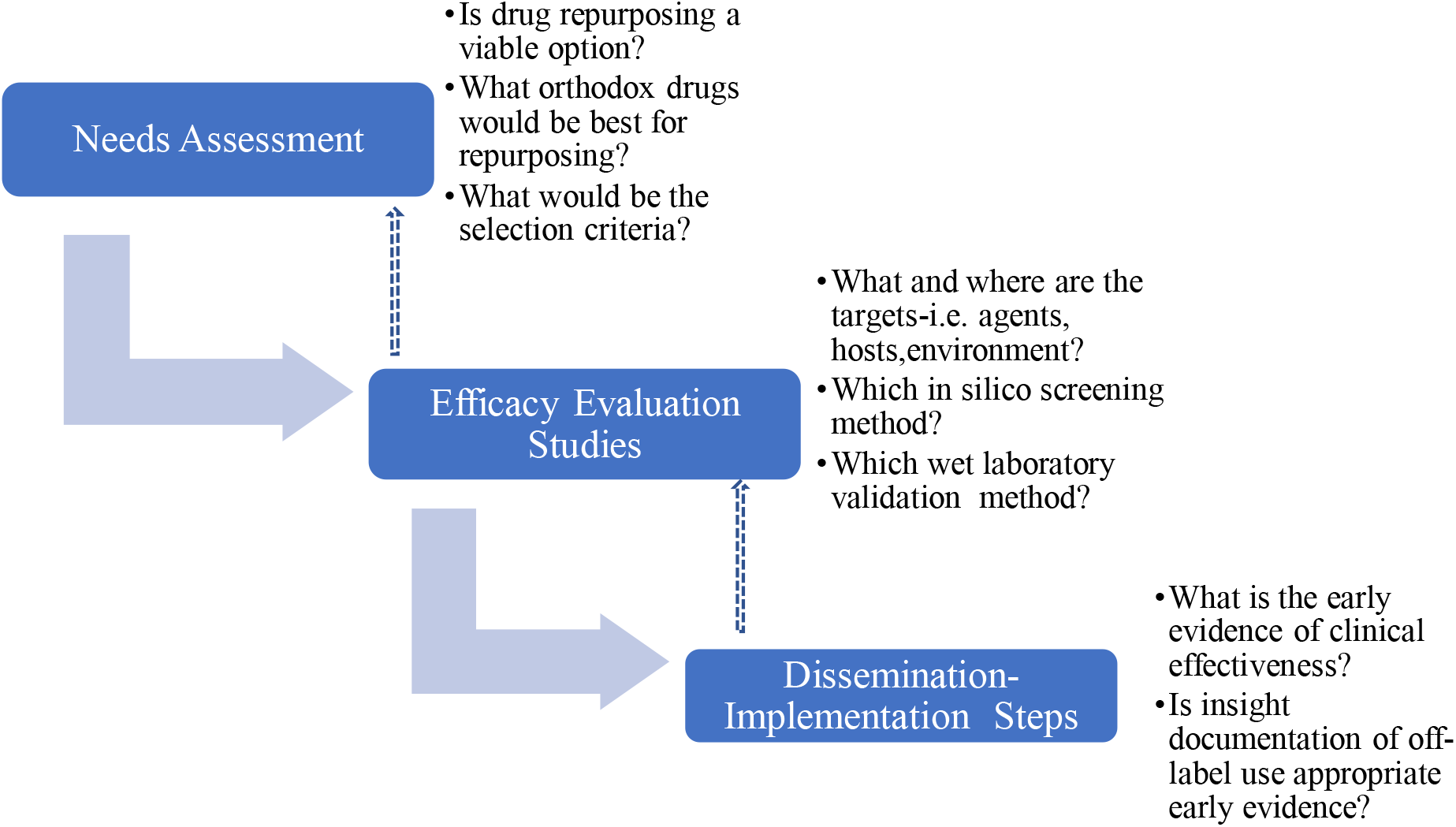
Schema of drug repurposing algorithm, with dissemination implementation design

Needs Assessment

#### 2.2.1 Needs Assessment

This involved developing a drug repurposing group that included those who will be directly involved in early phase laboratory studies (SOB, AY, AAA), later phase wet laboratory validation studies (SOB, AY, AAA, MUI, MBB, AS), clinical pharmacologist (SOB), policy makers(MAP,CLO), public health experts (EI,CLO, KE) and infectious disease clinicians (ZH) some of whom did not start the study but iteratively became involved as COVID-19 scenario evolved and needs became recognized. Consensus were reached using modified Delphi protocols involving relevant experts at each stage, with multiple iterations. The process involved questions and brainstorming on what drugs to select for repurposing efforts that will translate to feasible clinical implementation. In this regard, the opinions of infectious disease clinicians and clinical pharmacologists were considered particularly important because they may be the earliest advocates and champions of off-label use. For example, irrespective of potential for efficacy it will be inconceivable and a herculean task to attempt to repurpose digoxin for a viral infection because clinicians are unlikely to offer digoxin off-label irrespective of potential benefits^34,35^. The expected outcome of the needs assessment is to itemize a set of criteria for selecting drug candidates for repurposing.

#### 2.2.2 Efficacy Evaluation studies

##### 2.2.2.1 Selection of Drugs (ligands), SAR-CoV-2 protein (targets), and development of local database

The criteria for selecting drug candidates, previously developed, were applied to drugs with FDA-registration and whose structures was available on Drugbank(www.drugbank.com). One thousand four hundred and ninety drugs that met all the criteria were, therefore, downloaded.

Iterative scoping literature review was then conducted to understand the emerging knowledge of life cycle of SARS-CoV-2 virus as well as the host pathways involved in COVID-19. The review was conducted by a subgroup (SOB, AY, AAA) of the repurposing group and was continuously updated. The results were updated to all current members of the group in an iterative fashion.

The crystal structures of SARS-CoV-2 proteins critical in the life cycle of the virus, as well as their critical role in the pathogenesis of Covid-19, were identified based on a literature search and downloaded from the Protein Data Bank(www.rcsb.org) provided they also met the criteria as highlighted by Warren et al^36^ for selection of crystal structure for molecular docking, namely availability of experimental data for the protein, an R-free value of less than 0.45, the difference between R and R-free values should be equal to or less than 0.05, a density precision index of < 0.5, and structure with a higher resolution of less than 3.5Å. A local database of both the ligands and targets was then created on a desktop computer.

##### 2.2.2.2 Molecular Docking Using Schrodinger Maestro

The computer simulations were run on a 27 inches iMac desktop workstation using the Maestro graphical user interface of industry standard Schrodinger software (www.schrodinger.com). Hardware specifications include Graphics-AMD Radeon Pro 5700 8 GB, Memory-24 GB 2133 MHz DDR4, Processor-3.6 GHz 10-Core Intel Core i9 and 5K retina display all of which were optimized for structural visualization. All the molecular dockings were performed using Maestro Schrodinger version 9.2 on default settings and using protocols previously established in several studies^37–40^.

###### 2.2.2.2.1 Protein Preparation, SiteMap, and Grid Box generation

SARS-CoV-2 proteins (targets) stored in the local database were uploaded onto Maestro software and prepared using the Protein Preparation Wizard, keeping the settings at default. ‘Full’ protein preparation was done, which includes both H-bond network optimization and geometry minimization, as these have been shown to give significantly better results than the minimal preparation^41,42^. Briefly, the structures were loaded into the Maestro workspace by clicking the Import and Process tab, of the wizard. Thereafter, unwanted chains and waters beyond 5Å away from the binding pocket and Het groups were deleted by clicking the Review and Modify tab. Finally, the orientations of hydrogen-bonded groups and energy of the structures were optimized and minimised using optimized potentials for liquid simulations 2005 (OPLS-2005) force field by clicking the Refine tab. To check for errors due to automation, the prepared structures were reviewed manually for residues that have missing atoms, overlapping atoms, and atoms that are incorrectly typed. Also, a Ramachandran Plot was developed and inspected for stearic clashes.

The location of the primary binding site on the prepared proteins was predicted using the SiteMap module of the Schrödinger software suite. SiteMap uses the interaction energies to find the most energetic favourable areas. First, it traces the sites that include multiple site points on a grid. The number of site points for a site was set to 15 and the number of sites to be found was set to 5 and a more restrictive definition of hydrophobicity and OPLS force field were used for all the calculations^43^. Binding pockets site-mapping is a robust method of identifying more potential binding sites than limiting to the binding pocket of the co-crystallized ligand^41^. More so, in some cases, the location of a binding site for protein-ligand or protein-protein interactions is not resolved even though the protein structures are available^42^.

Furthermore, receptor grid boxes were generated using Glide’s Receptor Grid Generation module. A binding site residue identified by the SiteMap tool was selected and served as the grid-defining area for all the respective targets. Default Van der Waals radius scaling parameters were used (scaling factor of 1, partial charge cut-off of 0.25) at the binding pocket with the radius of 20 Å around the sitemap, setting the dimension of the grid box at 10 Ǻ × 10 Ǻ × 10 Ǻ.

###### 2.2.2.2.2 Ligand preparation

LigPrep wizard from the Maestro builder panel was used to prepare ligands and generate a 3D structure of the ligands in the ionization/tautomeric states using Epik. Hydrogen atoms were added while salts were removed at ionizing pH (7 ± 2). Energy minimization was performed using OPLS-2005 force field by using the standard energy function of molecular mechanics and RMSD cut-off 0.01Ǻ to generate the low-energy ligand isomer. The top-ranking ligand conformation was selected for the subsequent molecular docking.

###### 2.2.2.2.3 In-silico ligand to target interaction studies(docking)

Once the protein and ligand were prepared and receptor grid-generated, ligands were then docked to the proteins using Grid-based Ligand Docking with Energetics (GLIDE) docking protocol. The ligands were docked using Extra Precision (XP) mode. The docked conformers were evaluated using the Glide (G) Score. The G Score was calculated as follows^44,45^:

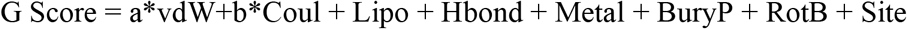

Where, vdW means Van der Waals energy, Coul means Coulomb energy, Lipo means lipophilic contact, HBond designates hydrogen-bonding, Metal indicates metal-binding, BuryP means penalty for buried polar groups, RotB indicates a penalty for freezing rotatable bonds, Site denotes polar interactions in the active site and the a=0.065 and b=0.130 are coefficients of vdW and Coul. The types of ligands-protein interactions were first evaluated and provided they form hydrophilic, hydrophobic, and electrostatic interactions, the ligands were then ranked from the arithmetic sum of the docking scores in the Standard Precision method (SP-docking).

#### 2.2.2.3 Wet Laboratory Validation of in-silico studies by inhibition of SARS-CoV-2 induced CPE

Wet laboratory evaluation of drug’s capacity to inhibit SARS-CoV-2 CPE was performed using the methods described and validated by Severson et al^20^.

Analytic standards of drugs that met the criteria for selection after the in-silico screening were purchased from MedChem Express (USA); all were purchased already dissolved in DSMO. Other materials used in this study include SARS-CoV-2 virus (P_3_), Glacial acetic acid, Ethanol 96% 0.4% (wt./vol), Trypan blue in 0.9% NaCl solution, Vero cells, SARS-CoV-2 clinical sample (ID 1239), Minimal Essential Medium/Earls Balance Salts (MEM/EBSS) (HyClone laboratories, Utah, USA), Trypsin, Tissue cultureware, fetal bovine serum (FBS) (Gibco, Life Technologies Ltd., Paisley, UK), and Penicillin/streptomycin (PenStrep).

##### 2.2.2.3.1 Culturing SARS-CoV-2 in Vero-6 cells

Vero cells were thawed and transferred into a 15-mL conical tube containing 10mL of MEM/EBSS supplemented with 10% FBS and 1% Penstrep (this step was performed to remove the cryo-preservative-dimethyl sulfoxide (DMSO)). The 15 mL tube was then centrifuged at room temperature for 5 minutes at 200 g. The supernatant was discarded, and the cells were resuspended in 5 mL of MEM/EBSS containing 10% FBS and 1% Penstrep. The Vero cell suspension was transferred to a 25 cm^2^ (T25) tissue culture flask with a vented cap and incubated at 37 °C in 5% CO2. The cells were monitored daily, and the media was changed every 3 days.

At about 90% confluent monolayer, the cells were subcultured into a new tissue culture flask. Confluent monolayer of Vero cells was detached using 0.25% trypsin-EDTA (Gibco) and resuspended in MEM/EBSS containing 10 % FBS and with the use of hemocytometer and trypan blue exclusion method; about 2.1 million Vero cells were counted and seeded in a T-75 flask containing MEM/EBSS plus 1% FBS without Penstrep and incubated for about 24 hours at 37°C in 5% CO_2_. Thereafter the flask was viewed under an inverted microscope for confluence. A confluence of 70 to 80% was desirable. Meanwhile, a positive SARS-COV-2 sample (sample ID 1239) confirmed by PCR, with a Circle threshold (CT) value of 12, obtained from CAMRET Sokoto, was filtered with a 0.22 µm Membrane filter. Once the 70 to 80% confluence monolayer was achieved, the medium was removed leaving about 2.5 ml. Then, 250 µl of the filtered virus sample was added to the flask and incubated for 1 hour at 37°C in 5% CO_2_. Thereafter, the media was replenished up to a total volume of 10 ml with MEM/EBSS having 1% FBS without Penstrep. The flask was incubated for 96 hours at 37°C in 5% CO_2_. CPE was checked daily with an inverted microscope until significant CPE was seen. The supernatant was then collected from the infected flask and centrifuged for 5 min at 500 × g, room temperature, to remove any cellular debris. The clarified supernatant was then stored in 1.5-ml screw-cap tubes at -80°C in aliquots of 1 ml until use.

##### 2.2.2.3.2 Inhibition of SARS-CoV-2 induced CPE by selected drugs-*rapid qualitative assay*

This study was designed as a high-throughput qualitative assessment of CPE inhibition by candidate drugs. Drugs that show positive CPE inhibition will then be tested in dose ranges in a separate study; thus, reducing the cost and time of this initial screening. All candidate drugs were screened at the maximum achievable plasma concentration (C_max_) at the recommended dose, as identified in literature. This concentration will also be the maximum tested in subsequent experiments to avoid spurious efficacy at concentrations (in vitro) not achievable in vivo. Such non-physiological result was a major drawback of several Ivermectin repurposing studies. The controls were three, as follows: (1) vehicle control groups (DMSO), (2) a negative control (Vero cells infected with SARS-CoV-2 virus without any treatment) and (3) a Vero cells growth control. All experiments were run in triplicates.

Briefly, we seeded 96-well plates with 6x10^4^ cells/mL of Vero E6 (200 μL per well), using MEM with 10% of fetal bovine serum (FBS) without antibiotics. Plates were incubated overnight at 37 °C in a 5% CO2 atmosphere. The following day, the 96 wells plates were viewed under an inverted microscope for confluence of about 50%.

Before drug treatment, cell culture supernatant was removed from each well and the wells were washed with 150 μL phosphate buffer solution (PBS). Except for the negative and cell growth control wells where only PBS (50 μL) was added, each well was infected with 50 μL SARS-CoV-2 diluted in PBS at a multiplicity of infection (MOI) of 0.1. All were then incubated for 1 hour at 37 °C in 5% CO2 with intermittent shaking of the plates at 15 minutes interval to allow for viral adsorption. Thereafter, the supernatant was removed and 200 μL of the respective drugs diluted in MEM having 1% FBS without antibiotics were added to the different treatment groups and incubated at 37 °C in 5% CO2. The cells were viewed using an inverted microscope at 48 hours to check for CPE. Two investigators (MBB and AAA) jointly visually assessed the slides and agreed on the percentage inhibition of CPE demonstrated by the test drugs.

## 3. Results

### 3.1 Outcome of Needs Assessment

#### 3.1.1 Selection criteria and rationale -Drug targets

Thirteen drug targets selected after several iterations of modified Delphi protocol, extending to the end of wet laboratory studies, are shown in Table 1 and Figure 2. Major targets are those assessed as critical for SARS-CoV-2 replication and are non-redundant because there are no alternative pathways to replication that exclude these enzymes. All other targets were classified as minor for the contrary reasons. Targets are considered validated if at least one publication exists that demonstrated that inhibition of the target resulted in reduction of viral replication.

**Table 1:** Drug targets and rationale*

| S/N | Name | PDB-ID | Target classification | Rationale |
| --- | --- | --- | --- | --- |
| 1 | SARS-CoV-2 RNA-dependent **RNA polymerase** in complex with cofactors | **6M71** | Major | A critical enzyme in viral replication and has no redundant pathway. It has been validated as a drug target<sup>61,62</sup> |
| 2 | The crystal structure of SARS-CoV-2 **main protease** in complex with an inhibitor N3 | **6LU7** | Major | A critical enzyme in viral replication. It has been validated as a drug target<sup>63</sup> |
| 3 | The crystal structure of SARS-CoV-2 **main protease** in apo form | **6M03** | Major | Critical enzyme and validated<sup>63</sup> |
| 4 | Structure of SARS-CoV-2 **main protease** bound to potent broad-spectrum non-covalent inhibitor X77 | **6W63** | Major | Critical enzyme and validated<sup>63</sup> |
| 5 | Structure of post fusion core of SARS-CoV-2 S2 subunit (fusion of the viral and cellular membranes)-**spike** | **6LXT** | Minor | Important entry protein, validated but redundant<sup>64</sup> |
| 6 | Structure of the SARS-CoV-2 HR2 Domain of **spike**: heptad repeat 2 (HR2) - bringing viral and cellular membranes in proximity for fusion | **6LVN** | Minor | Important entry protein, validated but redundant<sup>64</sup> |
| 7 | Crystal structure of SARS-CoV-2 **receptor binding domain** in complex with human antibody CR3022-**Spike** | **6W41** | Minor | Important entry protein, validated but redundant<sup>64</sup> |
| 8 | Prefusion SARS-CoV-2 **spike** glycoprotein with a single **receptor-binding domain**(RBD) up | **6VSB** | Minor | Important entry protein, validated but redundant<sup>64</sup> |
| 9 | The SARS-CoV-2 **RBD/ACE2-B0AT1** complex-**ACE2** | **6M17** | Minor | Important enzyme, validated but redundant<sup>65</sup> |
| 10 | Crystal structure of SARS-CoV-2 **nucleocapsid** protein N-terminal RNA binding | **6M3M** | Minor | Important viral protein and involved in COVID-19 pathogenesis, validated but redundant<sup>66,67,68</sup> |
| 11 | Crystal structure of the hexameric human **IL-6/IL-6 alpha receptor**/gp130 complex | **1P9M** | Minor | Important receptor in COVID-19 pathogenesis, validated but redundant<sup>69,70</sup> |
| 12 | Structure of the human **NF-kappaB p52 homodimer-DNA complex** at 2.1 Å resolution | **1A3Q** | Minor | Important factor in COVID-19 pathogenesis, validated but redundant<sup>70,71</sup> |
| 13 | The crystal structure of **Papain-Like Protease of SARS CoV-2** in complex with PLP_Snyder530 inhibitor | **7JIW** | Minor | Important enzyme, validated but redundant<sup>72,73</sup> |

\*Validated SARS-COV-2 and COVID-19 druggable targets: (6VSB, 6W41, 6LVN, 6LXT, 6M17, 6M71, 6W4B, 6LU7, 6W63, 6M03, 6M3M, 1P9M and 1A3Q)<sup>68,74–82</sup>

**Figure 2:**
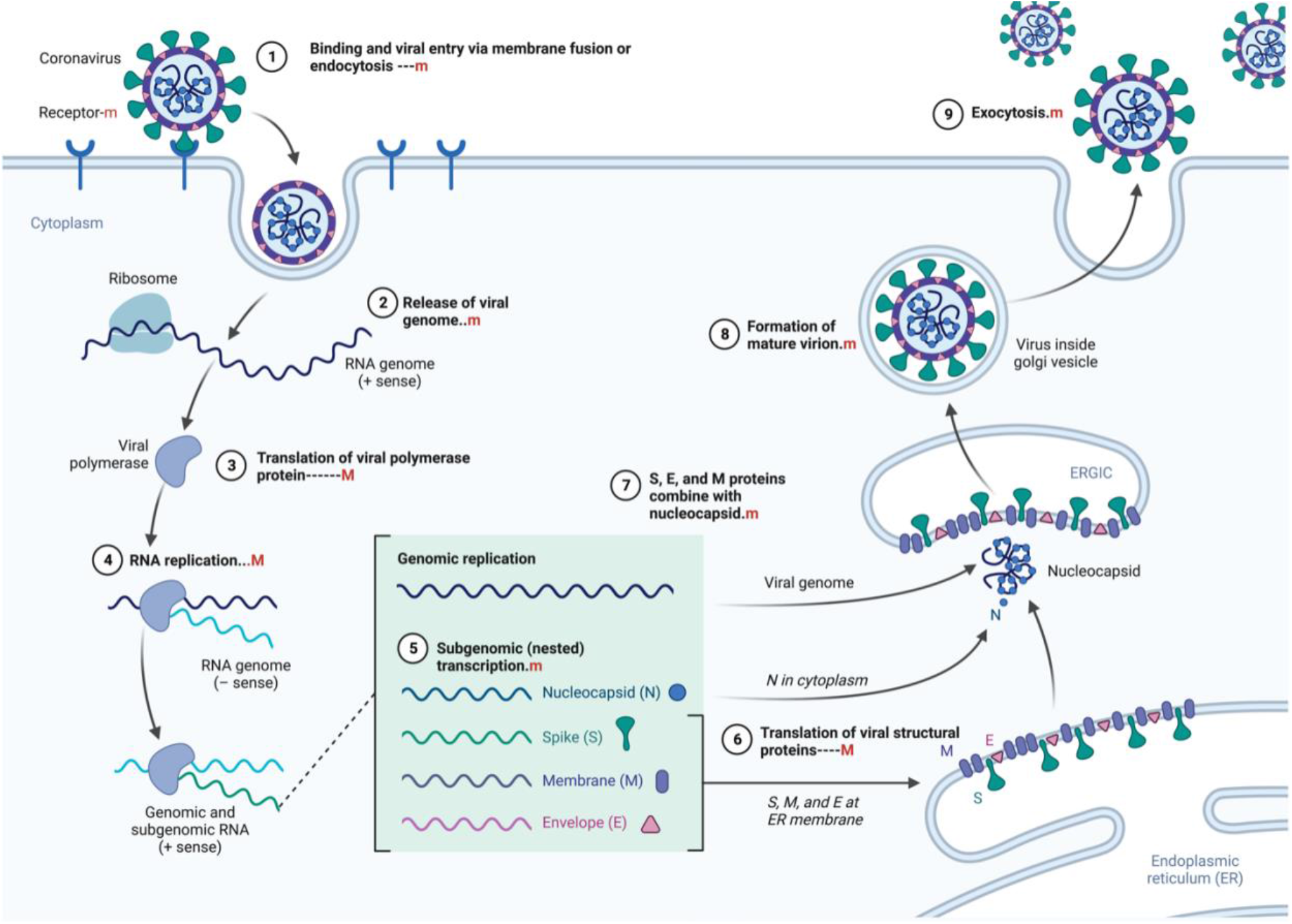
Replication cycle of SARS-CoV-2 and drug targets; ‘M’= major Targets and ‘m’= minor target

#### 3.1.2 Selection criteria and rationale -Drugs for repurposing for COVID-19

Five criteria developed for selecting drugs for repurposing after modified Delphi protocol iterations extending to the end of wet laboratory studies are shown in Table 2. All criteria must be met for a drug to be selected. The group of experts expect that drugs that meet these criteria should be rapidly available in all parts of the world if the drugs are found to be effective and will have a high threshold before resistance would emerge.

**Table 2:** Criteria for selecting drugs for repurposing and rationale for the selection

| S/N | Criteria | Rationale |
| --- | --- | --- |
| 1 | Negative binding energy to at least 3 targets, two of which must include SARS-CoV-2 RNA-dependent RNA polymerase and SARS-CoV-2 Main Protease (M<sup>Pro</sup>) enzyme | This is expected to reduce the chance of viral resistance to above 1 in 10<sup>*18</sup> |
| 2 | Provided criteria 1 is met, lowest overall bindings energies summed across all targets is preferred | This is expected to further reduce the probability of viral resistance and it also includes non-viral targets which increase the probability of therapeutic success |
| 3 | Provided criteria 1 and 2 are met, drugs whose patents have expired are preferred | This is expected to enable production of generic version and therefore increase availability. |
| 4 | Provided criteria 1 and 2 are met, drugs with high therapeutic indices are preferred. | This is expected to enhance the probability of off-label use from the patient and provider' perspective . |
| 5 | Provided criteria 1 , 2 and 4 are met, antimicrobials are preferred. | Antimicrobials are usually selectively toxic and have higher user (provider and patient) experience and therefore more likely to be prescribed off-label. |

The rationales are expected outcomes based on consensus between subject experts in a modified Delphi process.

### 3.2 Outcome of molecular docking and final selection for wet laboratory screening

Ninety-four drugs met the criteria for selection based on the docking result, Figure 3 and supplementary 1. Of these ninety-four, Streptomycin had the most favorable sum of scores across the targets (-109.1) while Tolbutamide had the least favorable docking across targets (-16.8). Nonetheless, after applying other criteria, Table 3 were the drugs selected for wet laboratory validation based on ability to inhibit SARS-CoV-2 induced CPE in Vero cells.

**Table 3:** Drugs selected for wet laboratory studies and rationale

| S/N | Drug | Rationale | Concentration tested | Rationale for Choice of Screening Concentration |
| --- | --- | --- | --- | --- |
| 1. | Erythromycin | Met all 5 selection criteria | 5µM | C<sub>max</sub> of after 500mg oral dose of erythromycin<sup>83</sup> |
| 2. | Pyridoxine | Met all 5 selection criteria | 10µM | C<sub>max</sub> of after 100mg oral dose of Pyridoxine<sup>84,85</sup> |
| 3. | Retapamulin | Met 4 of 5 selection criteria but attractive as a hand cream | 5µM | Estimated from the concentration in the 1 % ointment<sup>86</sup> and the Minimal Inhibitory concentration<sup>87</sup>. |
| 4. | Folic acid | Met all 5 selection criteria | 5µM | C<sub>max</sub> of after 400µg oral dose of Folic acid<sup>88</sup> |
| 5. | Ivermectin | Met all 4 of 5 selection criteria but attractive as a positive control | 5µM | Reflects maximum concentration tested and confirmed as effective invitro against SARS-CoV-2 from several studies<sup>18,89,90</sup> |

**Figure 3:**
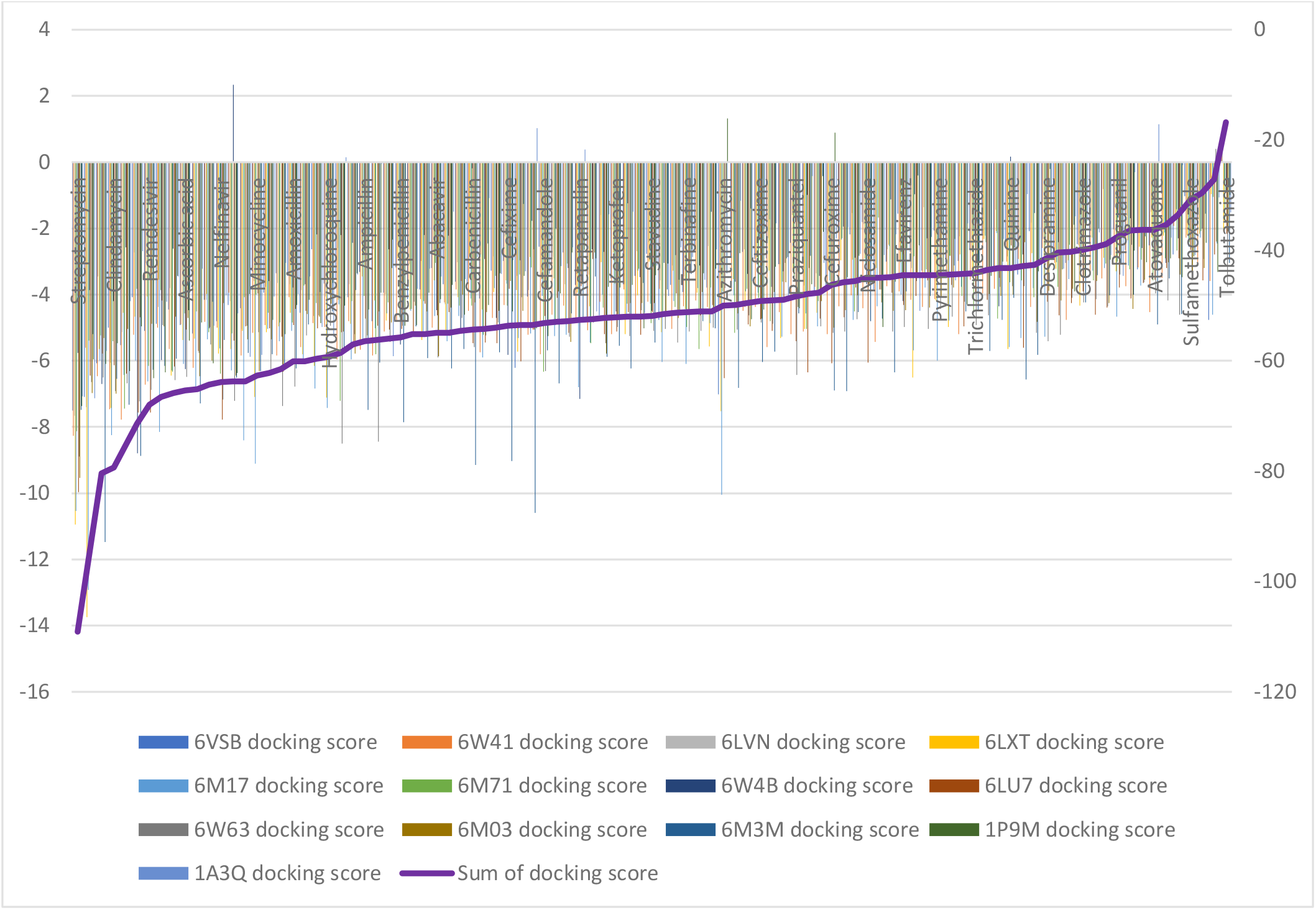
Docking scores of ninety-four drugs with favorable binding energies at all targets Drugs are ordered from left to right in descending favorable score. Only every 3^rd^ drug is shown in bold. The primary value axis(left) shows the docking score at each target while the secondary value axis(right) shows the docking score summed across all targets. The purple line across the figure represents the trend of the sum of docking scores. Streptomycin and Clindamycin have very favorable scores, but their nephrotoxicity and ototoxicity would reduce their chance of moving forward for repurposing in this schema given that alternatives exist, e.g., Ascorbic acid would be more favored.

### 3.3 Inhibition of SARS-CoV-2 induced CPE in Vero Cells

In this rapid qualitative assay, at the concentrations tested (C_max_ of the drugs), all the drugs inhibited SARS-CoV-2 induced CPE in Vero cells (Figure 4) with percentage inhibition ranging from 60%-80%, when compared to SARS-CoV-2 infected controls. Erythromycin caused the highest inhibition of CPE compared to other drugs tested, all at their C_max_ routine dose schedules.

**Figure 4*:**
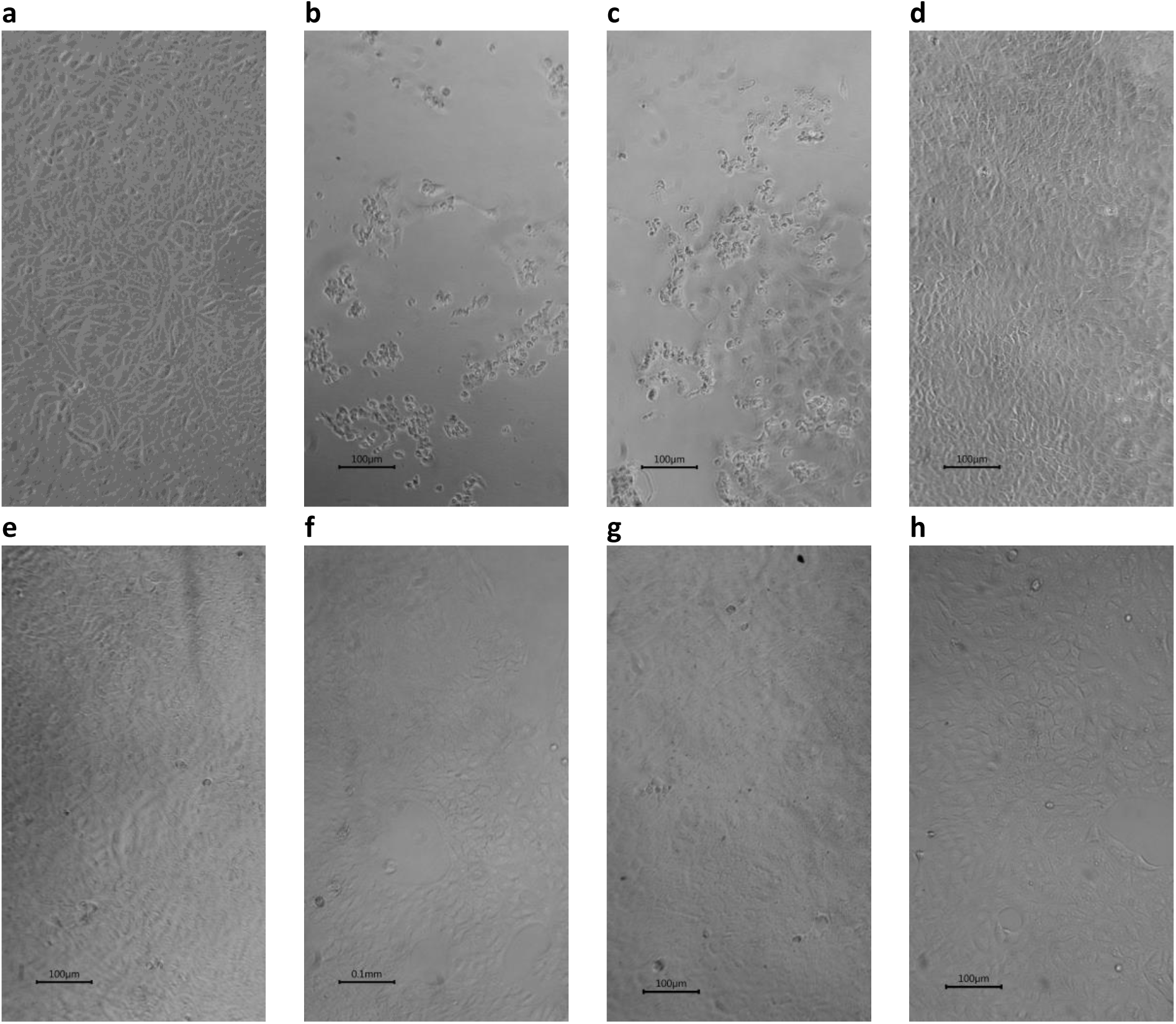
Inhibition by selected drugs of SARS-CoV-2 induced CPE in Vero cells(x10) **a**. Vero cells **b**. SARS-CoV-2 infected Vero showing ≥ 80% Cytopathic effects (CPE) **c**. SARS-CoV-2 infected Vero cells with 0.1% DMSO showing = 80% CPE **d**. SARS-CoV-2 infected Vero cells with 5μM Erythromycin in 0.1% DMSO showing ≥ 80% inhibition of CPE **e**. SARS-CoV-2 infected Vero cells with 5 μM Retapamulin in 0.1% DMSO showing ≥ 60% inhibition of CPE. **f**. SARS-CoV-2 infected Vero cells with 10 μM Pyridoxine in 0.1% DMSO showing ≥ 60% inhibition of CPE. **g**. SARS-CoV-2 infected Vero cells with 5 μM Folic acid in 0.1% DMSO showing ≥ 60% inhibition of CPE **h**. SARS-CoV-2 infected Vero cells with 5 μM Ivermectin in 0.1% DMSO showing ≥ 80% inhibition of CPE. *All images were taken and prepared by AAA, then reviewed by MBB.

## 4. Discussion and Conclusion

In this study, a novel algorithm was developed and implemented for repurposing drugs for emerging viral diseases using SARS-CoV-2 and COVID-19 as case studies. At each step, scenarios sought to maximize the probability of identifying an effective agent that may easily translate to clinical application through off-label principles while awaiting further studies. Also, the protocol was designed to be easily executed even in LMIC using CPE. These objectives appear successful because erythromycin, pyridoxine and folic acid that this study found effective in inhibiting SARS-CoV-2 induced CPE are almost ubiquitously available in the world and are accepted to be of low safety concerns^46–49^, especially when life time dosing is not under consideration. Indeed, Folic acid has previously been suggested to be a potential inhibitor of SARS-CoV-2 M^Pro 50,51^and SARS-CoV-2 nucleocapsid protein^52^, but these studies stopped at in-silico docking and focused on a single target. In a separate study, folic acid was shown to inhibit SARS-CoV-2 invasion of cells by methylating ACE2 and reducing the transmissibility in mice^53^, altogether suggesting a multitarget effect. In COVID-19 patients, serum levels of folic acid have been correlated with clinical outcome ^54^and high dose folic acid has been suggested as possible add on therapy or supplement for COVID-19^55^. Paradoxically folic acid has been used as inert placebo in clinical trials of COVID-19 infection^56^ instead of as an active comparator thereby possibly biasing the result. Supplements,with pyridoxine 4.9 g daily and folic acid 400μg daily, as a component, have been shown to improve COVID-19 clinical outcome^57^. Erythromycin, the prototype macrolide antibiotic, has not been much studied for SARS-CoV-2 except in computational docking ^58,59^, which suggested activity against SARS-CoV-2 M^Pro^, but these studies also mainly have single target.

Retapamulin is less ubiquitous and will probably be difficult to access in LMIC, nevertheless, it is attractive as a potential hand or surface ointment solution. Ivermectin has previously been established to inhibit SARS-CoV-2 replication but at physiologically impossible dose^60^. In this study ivermectin served as a positive control and the result supports that inhibition of CPE is its additional or consequential effect and further reassures that the overall study design could be predictive. Use of qualitative assessment of CPE inhibition is one of the drawbacks of this study but it is conceivably more useful for rapid screening of large library of compounds compared to quantitative methods. Further studies of the drugs identified are needed and may include quantification of CPE inhibition, graded dose response of CPE inhibition, insight into mechanisms of actions and clinical trials. Dose response studies would be particularly important if doses beyond posology are under consideration; a likely situation where current clinical doses are not expected to be effective (like ivermectin) or have been shown not to be effective in previous clinical trials. Meanwhile, we propose that the result of the present study should be sufficient for off-label prescription of erythromycin, pyridoxine, and folic acid in COVID-19, and is particularly important during rapidly evolving, high mortality epidemics.

In conclusion, drug repurposing efforts need to be purposeful to optimize yields and avoid identifying drugs whose translation to clinical use may be compromised due to toxicity, availability, or inappropriate dosing requirements. This study has presented results that suggest that erythromycin, pyridoxine, and folic acid could be useful in the treatment of COVID-19.

